## Supplemental Table S1 for "Replication stress inducing ELF3 upregulation promotes BRCA1-deficient breast tumorigenesis in luminal progenitors"

### Supplemental Tables

Table S2.Detailed information on the BRCA1 germline mutation and clinicopathological characteristics of seven patients.

| Patient ID | BRCA germline mutations | age(y) | parity | pathologic | ER | PR | HER2 | HER2 FISH | Ki-67 | PAM50 subtype | neoadjuvant therapy |
|---|---|---|---|---|---|---|---|---|---|---|---|
| Case_1 | c.4013delA (p.Lys1338Argfs*28) | 25 | nulliparous | IDC | 90%+ | negative | 2+ | negative | 20%+ | Luminal A | none |
| Case_2 | c.2194G>T (p.Glu732Ter) | 43 | multiparous | IMC | 20%+ | negative | 2+ | negative | 70%+ | Luminal B | chemotheray |
| Case_3 | c.1069A>T (p.Lys357Ter) | 29 | multiparous | IDC | negative | negative | 2+ | negative | 80%+ | Basal-like | none |
| Case_4 | c.5251C>T (p.Arg1751Ter) | 43 | multiparous | IDC | negative | negative | 1+ | NA | 60%+ | Basal-like | none |
| Case_5 | wild-type | 42 | multiparous | IDC | 80%+ | 80%+ | 1+ | NA | 30%+ | Luminal A | none |
| Case_6 | wild-type | 29 | nulliparous | fibroadenoma | — | — | — | — | — | — | — |
| Case_7 | wild-type | 33 | nulliparous | fibroadenoma | — | — | — | — | — | — | — |

BRCA1 transcript: NM_007294.3

Abbreviations: ER, estrogen receptor; PR, progesterone receptor; HER2, human epidemal growth factor receptor 2; IDC, invasive ductal carcinoma; IMC,invasive micropapillary carcinoma; FISH, fluorescence in situ hybridization
