## Supplemental Table S2 for "Replication stress inducing ELF3 upregulation promotes BRCA1-deficient breast tumorigenesis in luminal progenitors"

Table S3. Reagents used in this study.

| Reagent | Source | Cat# |
|---|---|---|
| Antibodies | | |
| Anti-BRCA1 | Merck-Millipore | OP92 |
| Anti-ELF3 (Immunoblot) | abcam | ab133621 |
| Anti-ELF3 (Immunohistochemistry) | Sigma-Aldrich | HPA003479 |
| Anti-Chk1 | Santa Cruz | sc-8408 |
| Anti-Chk2 | Santa Cruz | sc-17747 |
| Anti-phospho-Chk1 (S345) | CST | 2348S |
| Anti-phospho-Chk2 (T68) | CST | 2661S |
| Anti-c-Myc | Beijing Biodragon | B1002 |
| Anti-γH2AX | CST | 9718S |
| Anti-53BP1 | Merck-Millipore | MAB3802 |
| Anti-BrdU | BD | BD345780 |
| Anti-BrdU | abcam | ab6326 |
| Anti-Ku80 | Wang lab | |
| Anti-β-actin | Beijing RIBIO | 1015t |
| Anti-GAPDH | ABclonal | AC033 |
| Chemicals | | |
| Doxycycline hydrochloride（DOX） | HARVEYBIO | D31646 |
| Hydroxyurea (HU) | HARVEYBIO | H31749 |
| Aphidicolin (APH) | Cayman | 14007 |
| Olaparib | Selleck | S1060 |
| Bleomycin (BLM) | Cayman | 19692 |
| Mitomycin C (MMC) | Coolaber | COL-CM7391 |
| Cisplatin | HARVEYBIO | HZB0054 |
| Cell Culture and Transfection | | |
| RPMI 1640 | HyClone | SH30809.01 |
| DMEM | Gibco | C11995500BT |

| | | |
|---|---|---|
| DMEM/F12 | Gibco | C11330500BT |
| FBS | Yeasen | |
| Tet system approved certified FBS | BI | 04-005-1A |
| Horse serum | Beijing ZOMANBIO | ZX110 |
| EGF | Macgene | CC102 |
| Insulin | Macgene | CC101 |
| Hydrocortisone | Macgene | CC103 |
| Cholera toxin | Macgene | CC104 |
| Polybrene | Yeasen | 40804ES86 |
| Liposomal Transfection Reagent | Yeasen | 40802ES03 |
| jetOPTIMUS | Polyplus | 117-01 |
| Lipofectamine RNAiMAX | Invitrogen | 13778150 |
| Gen OPTI-MEM | Macgene | CT007 |

Table S4 shRNA sequences used in this study

| **Name** | **Sequence (5'-3')** |
|---|---|
| shBRCA1 | CAGCTACCCTTCCATCATA |
| shELF3 | GCCGATGACTTGGTACTGA |
| shCtrl | |

Table S5 PCR primers used in this study

| **Name** | **Sequence (5'-3')** |
|---|---|
| Human *ELF3* forward | ATGGCTGCAACCTGTGAGATTAGCA |
| Human *ELF3* reverse | TCAGTTCCGACTCTGGAGAACCTCT |
| Human *E2F6* forward | ATGAGTCAGCAGCGGCCG |
| Human *E2F6* reverse | TCAGTTGCTTACTTCAAGCAATTCTTC |

Table S6 siRNA sequences used in this study

| **Name** | **Sequence (5'-3')** |
|---|---|
| Negative control siRNA sense | CGUACGCGGAAUACUUCGATT |

| | |
|---|---|
| Negative control siRNA anti-sense | UCGAAGUAUUCCGCGUACGTT |
| Human *BRCA1* siRNA sense | CAGCUACCCUUCCAUCAUATT |
| Human *BRCA1* siRNA anti-sense | UAUGAUGGAAGGGUAGCUGTT |
| Human *ELF3* siRNA#3 sense | GAAGUGACGUGGACCUGGATT |
| Human *ELF3* siRNA#3 anti-sense | UCCAGGUCCACGUCACUUCCA |
| Human *ELF3* siRNA#4 sense | GCCGAUGACUUGGUACUGATT |
| Human *ELF3* siRNA#4 anti-sense | UCAGUACCAAGUCAUCGGCCC |
| Human *GATA3* siRNA sense | GCCUAAACGCGAUGGAUAUTT |
| Human *GATA3* siRNA anti-sense | AUAUCCAUCGCGUUUAGGCUU |

Table S7 RT-qPCR primers used in this study

| Name | Sequence (5’-3’) |
|---|---|
| Human *GAPDH* qPCR forward | CAACTACATGGTTTACATGTTC |
| Human *GAPDH* qPCR reverse | GCCAGTGGACTCCACGAC |
| Human *BRCA1* qPCR forward | CAACATGCCCACAGATCAAC |
| Human *BRCA1* qPCR reverse | ATGGAAGCCATTGTCCTCTG |
| Human *ELF3* qPCR forward | GTTCATCCGGGACATCCTC |
| Human *ELF3* qPCR reverse | GCTCAGCTTCTCGTAGGTC |
| Human *GATA3* qPCR forward | ACCACAACCACACTCTGGAGGA |
| Human *GATA3* qPCR reverse | TCGGTTTCTGGTCTGGATGCCT |
| Human *PRIM1* qPCR forward | TATCGCTGGCTCAACTACGGTG |
| Human *PRIM1* qPCR reverse | CACTCTGGTTGTTGAAGGATTGG |
| Human *PRIM2* qPCR forward | CTTCAGCCTCTGCTCAATCACC |
| Human *PRIM2* qPCR reverse | GTAACTGACGCATGCAAGGTGG |
| Human *PCNA* qPCR forward | CAAGTAATGTCGATAAAGAGGAGG |
| Human *PCNA* qPCR reverse | GTGTCACCGTTGAAGAGAGTGG |
| Human *MCM3* qPCR forward | CGAGACCTAGAAAATGGCAGCC |
| Human *MCM3* qPCR reverse | GCAGTGCAAAGCACATACCGCA |
| Human *MCM4* qPCR forward | CTTGCTTCAGCCTTGGCTCCAA |
| Human *MCM4* qPCR reverse | GTCGCCACACAGCAAGATGTTG |
| Human *MCM6* qPCR forward | GACAACAGGAGAAGGGACCTCT |
| Human *MCM6* qPCR reverse | GGACGCTTTACCACTGGTGTAG |

| | |
|---|---|
| Human *MCM7* qPCR forward | GCCAAGTCTCAGCTCCTGTCAT |
| Human *MCM7* qPCR reverse | CCTCTAAGGTCAGTTCTCCACTC |
| Human *RAD51* qPCR forward | TCTCTGGCAGTGATGTCCTGGA |
| Human *RAD51* qPCR reverse | TAAAGGGCGGTGGCACTGTCTA |
| Human *CDC7* qPCR forward | GGAAAACTGCCAGTTCTTGCCC |
| Human *CDC7* qPCR reverse | GGCACTTTGTCAAGACCTCTGG |
| Human *CDC45* qPCR forward | TGGATGCTGTCCAAGGACCTGA |
| Human *CDC45* qPCR reverse | CAGGACACCAACATCAGTCACG |
| Human *GINS2* qPCR forward | AGCCAAACTCCGAGTGTCTGCT |
| Human *GINS2* qPCR reverse | CTTGTGTGAGGAAAGTCCCGCT |
| Human *GINS4* qPCR forward | CTGGAGAGCAAGCCTGAGATTG |
| Human *GINS4* qPCR reverse | GCAAGTAGCTGCTGAGGACGTA |
